# Efficient enzyme coupling algorithms identify functional pathways in genome-scale metabolic models

**DOI:** 10.1101/608430

**Authors:** Dikshant Pradhan, Jason A. Papin, Paul A. Jensen

## Abstract

Flux coupling identifies sets of reactions whose fluxes are “coupled" or correlated in genome-scale models. By identified sets of coupled reactions, modelers can 1.) reduce the dimensionality of genome-scale models, 2.) identify reactions that must be modulated together during metabolic engineering, and 3.) identify sets of important enzymes using high-throughput data. We present three computational tools to improve the efficiency, applicability, and biological interpretability of flux coupling analysis.

The first algorithm (cachedFCF) uses information from intermediate solutions to decrease the runtime of standard flux coupling methods by 10-100 fold. Importantly, cachedFCF makes no assumptions regarding the structure of the underlying model, allowing efficient flux coupling analysis of models with non-convex constraints.

We next developed a mathematical framework (FALCON) that incorporates enzyme activity as continuous variables in genome-scale models. Using data from gene expression and fitness assays, we verified that enzyme sets calculated directly from FALCON models are more functionally coherent than sets of enzymes collected from coupled reaction sets.

Finally, we present a method (delete-and-couple) for expanding enzyme sets to allow redundancies and branches in the associated metabolic pathways. The expanded enzyme sets align with known biological pathways and retain functional coherence. The expanded enzyme sets allow pathway-level analyses of genome-scale metabolic models.

Together, our algorithms extend flux coupling techniques to enzymatic networks and models with transcriptional regulation and other non-convex constraints. By expanding the efficiency and flexibility of flux coupling, we believe this popular technique will find new applications in metabolic engineering, microbial pathogenesis, and other fields that leverage network modeling.

## Introduction

Constraint-based modeling allows genome-scale networks to be simulated using minimal kinetic information [1]. Using tools from mathematical programming, modelers can identify fluxes [2], gene expression levels [3, 4], and metabolite concentrations [5, 6] that satisfy thermodynamic, stoichiometric, and mass-balance constraints. These methods enable researchers to explore the physiology of microbes with as little as a genome sequence [7].

Flux balance analysis (FBA) and other constraint-based modeling techniques assume a steady state (or quasi-steady state) [8]. No metabolites accumulate or deplete during steady state. Consequently, the fluxes of several reactions in the metabolic model are *coupled*, or perfectly multicollinear [9]. Specifying the flux of any one reaction in a set of coupled reactions constrains the fluxes of the other reactions in the coupled set to a scalar multiple of the specified flux. The simplest example of a coupled set of reactions is a linear, unbranched pathway [8].

Coupled reactions are useful in at least three ways. First, in metabolic engineering, sets of coupled reactions can be modulated together to increase flux through a pathway, as coupled reaction sets may mimic natural regulatory patterns in microbes [10]. Second, coupled reaction sets can be used as surrogates for metabolic pathways. Genes associated with coupled reaction sets are enriched for genetic interactions and have correlated expression [11]. While “standard" pathway definitions exist for metabolic reactions, these definitions are biased toward the metabolic networks of model organisms like the bacterium *E. coli*, the yeast *S. cerevisiae*, or humans [12, 13]. Reactions in standard pathways may not function together in non-model organisms. Grouping reactions by function (e.g, as correlated reaction sets) may avoid the biases of pathway databases. Finally, finding coupled variables reduces the dimensionality of metabolic models. Since fully coupled variables are multicollinear, a set of coupled variables can be replaced with a single variable without loss of information. The reduced models may be more efficient when fitting parameters or applying computationally expensive algorithms [14, 15].

Applications of coupled reaction sets (like the examples above) focus primarily on the enzymes associated with the reactions, not the reactions themselves. While constraint-based algorithms often focus on fluxes, most experimental studies examine metabolic enzymes on the gene or protein level. Proteomics and especially transcriptomics data are easier and cheaper to acquire on a genome-wide scale than flux profiles, and many metabolic studies involve gene- or protein-centric views of metabolic pathways. Unfortunately, creating “coupled enzyme sets" from coupled reaction sets is not trivial [16]. Gene products associate combinatorially to catalyze reactions. Protein subunits can interchangeably combine to form enzyme complexes, so the relationship between gene products and the flux through reactions involves logical “or" and “and" operations. Enzymes can also be associated with multiple reactions across several pathways. The coupling between enzymes is different from the coupling between reactions due to the logic and promiscuity in the gene-protein-reaction (GPR) network. One cannot find coupled enzyme sets by collecting all enzymes associated with each coupled reaction set.

We present three algorithmic improvements for identifying coupled reactions and enzymes. First, we describe a mathematical transformation for constraint-based metabolic models that incorporates enzyme activities as continuous variables. We call this framework Flux and Activity Linked Constraints, or FALCON. For enzyme activities to be directly coupled to reaction fluxes, all models with reversible reactions must be described using a set of binary variables. The binary variables destroy the convexity of the FALCON model’s solution space, making the model more difficult to solve. Without a convex solution space, fast methods for identifying coupled reaction sets cannot be used. Instead, we present modifications to the original method for identifying coupled reactions (flux coupling finder, or FCF). By caching intermediate solutions, we should how an improved FCF algorithm can quickly find couplings between variables in convex and non-convex models.

Next, we compare coupled enzyme sets calculated directly from the FALCON model with sets found indirectly by mapping enzymes to coupled reaction sets. We show that when compared to enzymes associated with reaction sets, the FALCON-derived sets are smaller, have stronger correlation in expression levels, and are more likely to share fitness changes when deleted. Thus, the FALCON-derived sets behave in many ways as a single unit, consistent with our intuition regarding correlated sets.

Finally, we apply our faster flux coupling algorithm to identify sets of reactions or enzymes that include singly redundant pathways. By extending the definition of coupling, we can create enzyme sets from redundant enzymes that are widely considered to be in the same pathway but are not placed into coupled sets by previous methods. Enzymes in these expanded sets retain close ties regarding expression changes and phenotypic importance.

## Results and Methods

### Notation

We use the term “*R* sets” to denote sets of fully coupled reactions as defined by Burgard and Maranas [9]. A model’s *R* sets can be found using the Flux Coupling Finder (FCF) algorithm [9]. Our faster cachedFCF algorithm changes only the speed of finding *R* sets, not their definition. A standard metabolic model will produce the same *R* sets via FCF or cachedFCF.

The standard method to find enzyme sets is to collect all enzymes associated with the reactions in an *R* set using the gene/protein/reaction relationships in the model. Because these enzyme sets are constructed using the *R* sets, we call the corresponding enzyme sets “*E*_FCF_(*R*) sets”. Note that *R* sets are disjoint, i.e. each reaction in a model appears in exactly one *R* set. By contrast, the *E*_FCF_(*R*) sets are not disjoint (Figure 1B). Enzymes in a model can be associated with many reactions, and these many-to-many relationships can associate one enzyme with multiple *R* sets.

Finally, we refer to sets of enzymes calculated directly from a FALCON model as “*ɛ*_FALCON_ sets”. (Since *ɛ*_FALCON_ sets are not constructed from *R* sets, we do not write them as a function of *R*.) Unlike *E*_FCF_(*R*) sets, the *ɛ*_FALCON_ sets calculated from FALCON models are disjoint, regardless of the complexity of the enzyme associations in the model.

### Reactions and enzymes couple differently in the same model

Enzymes can be associated with multiple reactions, and vice versa. The many-to-many relationship between enzymes and reactions can be seen in the GPRs of constraint-based models. For example, a single enzyme in *P. aeruginosa* PAO1 is associated with 30 reactions (Figure 1a). By consequence, the mapping between enzymes and coupled reaction sets is also not one-to-one. The *P. aeruginosa* metabolic network contains 5 enzymes that each map to more than 10 difference sets of coupled reactions (Figure 1b). There are multiple methods for defining enzymes sets, even for small reaction networks. Consider the network of six reactions and seven enzymes shown in Figure 1c. Flux coupling identifies two coupled reaction sets (Figure 1d). We can create enzyme sets by mapping the reaction sets to their associated enzymes (Figure 1e). However, one enzyme appears in both sets even though the reaction sets are disjoint (Figure 1e). One of the enzyme sets also contains two isozymes (Figure 1e). It is unlikely that the isozymes are coupled (i.e. one has activity if and only if the other does), especially when many isozymes are not co-expressed [17]. A better partitioning of the enzymes appears in Figure 1f. The enzymes in each set are fully coupled. The enzyme sets in Figure 1f are the result of applying flux coupling to a metabolic network model that includes both reactions and enzymes as continuous variables using FALCON.

### Linking enzyme activity with reaction flux

Simulations on COBRA models with enzyme associations follow a two-step procedure [16]. The process begins with a genetic state describing the subset of genes in the model that are expressed for a set of model parameters (environment, mutations, etc.). In the first step, the rules for each reaction are evaluated in the context of the genetic state. A reaction is removed from the model if its rule is not satisfied. In the second phase, the sub-model containing all reactions with satisfied rules is solved or optimized. Separating GPR evaluation and flux optimization into two sequential stages prevents algorithms from fully interrogating the relationship between the enzymatic network and the reaction network. Multiple frameworks consolidate the GPR logic and reaction stoichiometry into a single optimization problem. One of the earliest frameworks, SR-FBA [19], treated enzymes as binary variables and added mixed integer constraints to enforce the GPR logic. Although SR-FBA includes both enzymes and reaction fluxes in a single optimization problem, one cannot identify fully coupled sets of enzymes when the enzymes are represented as binary, “on/off" variables.

To consider enzymes as continuous values in a model, multiple quasi-stoichiometric frameworks have been developed [20, 21]. These frameworks rewrite all reactions in a model to consume a pseudo-metabolite representing the activity of the associated enzyme (Figure 2). In the simplest case, an irreversible reaction *A* → *B* catalyzed by enzyme *e* is rewritten as *A* + Activity(*e*) → *B*, so every unit of flux through the reaction consumes not only a unit of the metabolite *A* but also a unit of activity for the enzyme *e* (Figure 2A). If two or more enzymes (or enzymatic subunits) are required to catalyze a reaction, a reaction in a FALCON model consumes activity from both enzymes (Figure 2B). When two or more enzymes can independently catalyze a reaction, activity from either enzyme can be consumed (Figure 2C). Separate reactions associated with the same enzyme all draw from the same pool of enzyme activity (Figure 2D). Sharing enzyme activity across reactions adds additional constraints from the enzyme association network that are not enforced when only reaction fluxes are included in a model.

**Figure 1.**
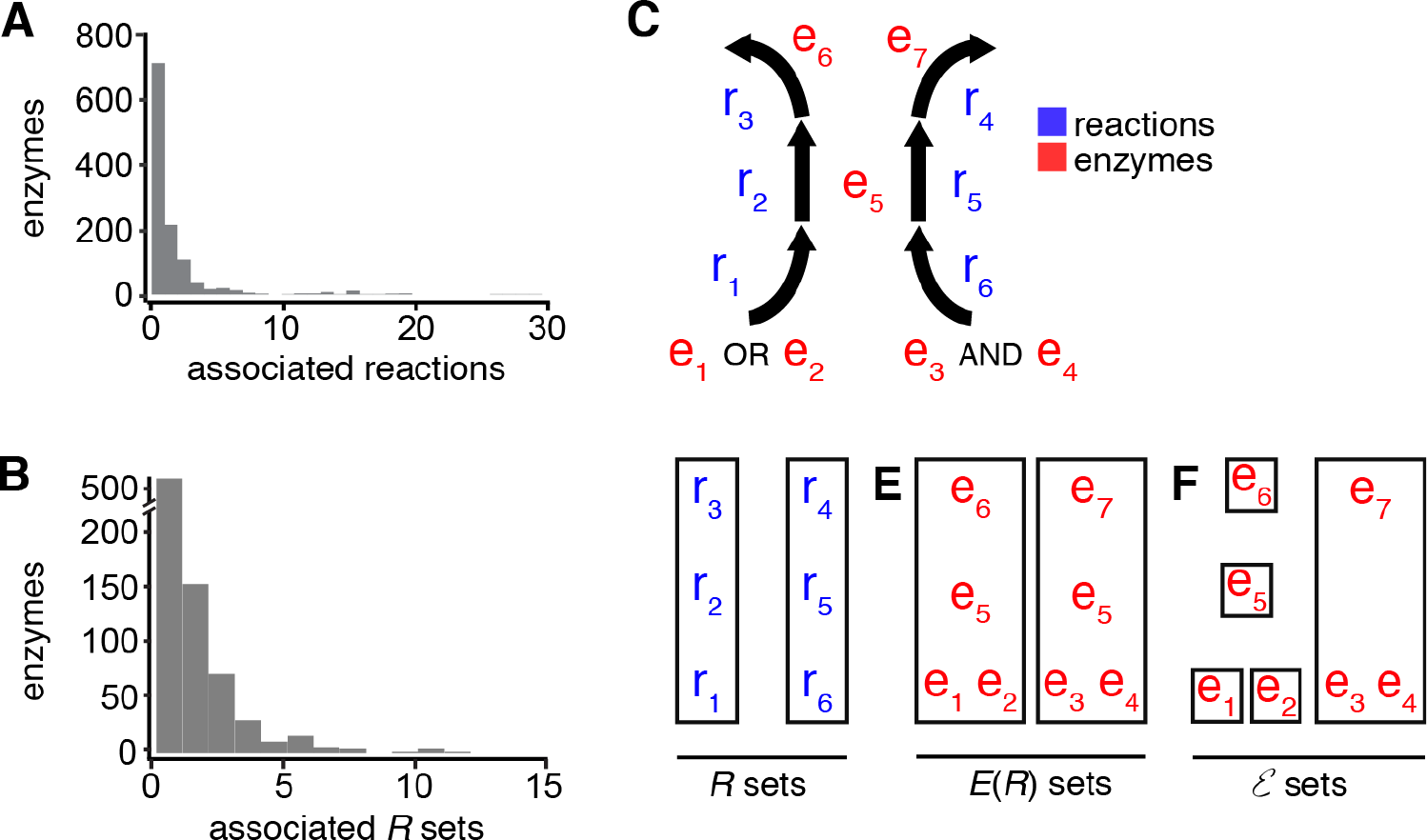
**A.** Enzyme associations in the *P. aeruginosa* metabolic model [18] are not one-to-one between enzymes and reactions. Counting the number of enzymes (vertical) used in the GPR of each reaction (horizontal) reveals that 38.9% of enzymes associate with multiple reactions. **B.** Enzymes are not uniquely associated with coupled reaction sets. Coupled reaction sets were identified with cachedFCF and enzymes were mapped using GPRs in the *P. aeruginosa* model. Many enzymes are associated with reactions that belong to separate coupled reaction sets. **C.** A small reaction network illustrates how complex enzyme associations and shared enzymes affect reaction coupling. **D.** If enzymes are ignored, the network contains two sets of coupled reactions. **E.** Simply mapping all associated enzymes onto the reaction sets produces two sets of enzymes. The enzyme sets are not disjoint (*e*_5_ is shared) and contain several enzymes that should not be coupled (*e*_1_, *e*_2_, *e*_5_, and *e*_6_). **F.** Calculating enzyme sets directly with a FALCON model identifies the correct couplings.

**Figure 2.**
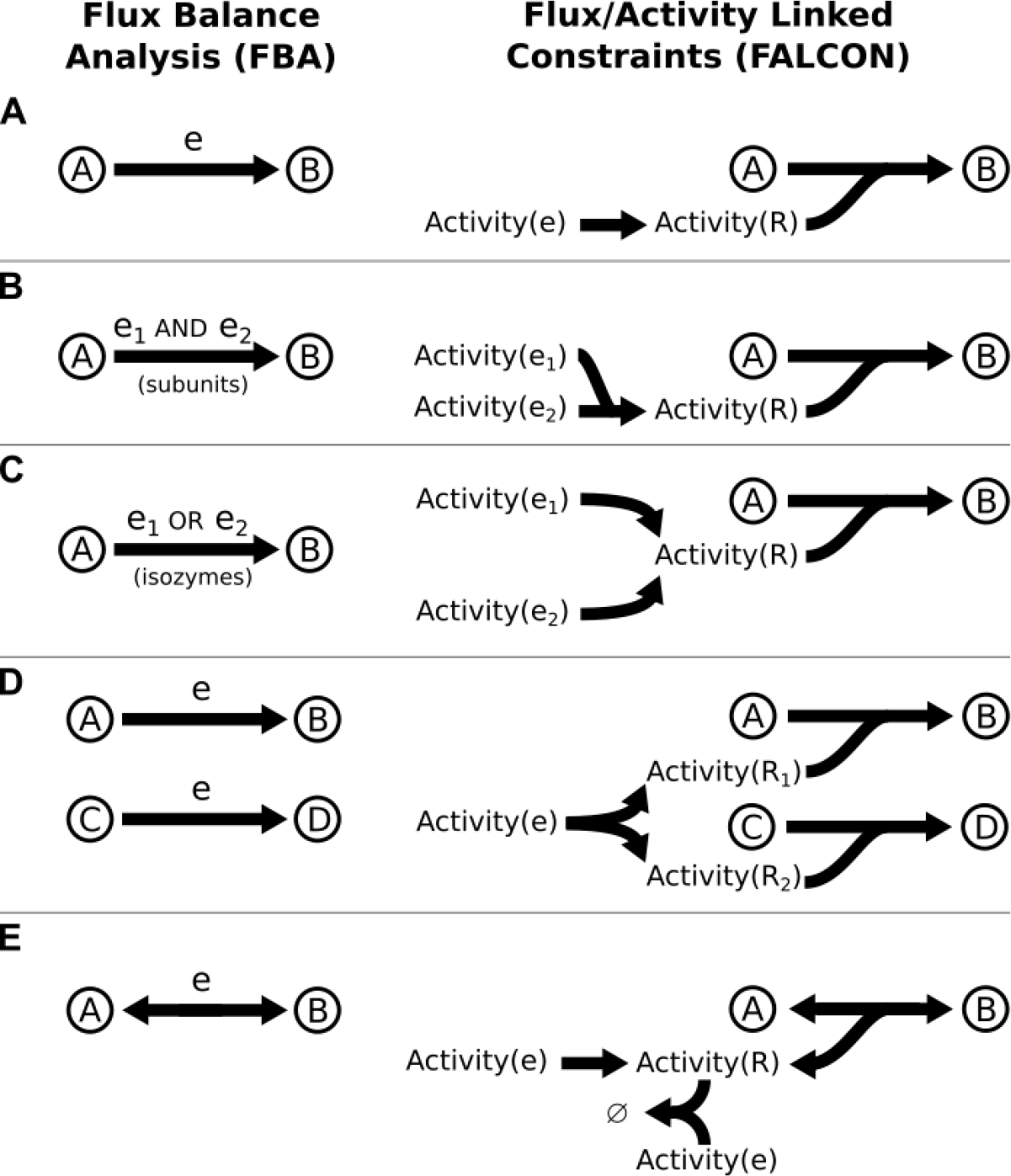
FALCON links enzymes with reactions by consuming a “pseudometabolite” representing enzyme activity (**A**). FALCON can represent enzymatic subunits (**B**), isozymes (**C**), and enzymes that are associated with multiple reactions (**D**). To avoid activity cycles, FALCON uses mixed-integer constraints to limit flux to a single direction in reversible reactions (**E**). Without the integer constraints, enzyme activity for reversible reactions could exceed the net reaction flux by an arbitrary value.

Our FALCON method can be applied to all enzyme associations, regardless of their complexity. The Boolean rule describing a reaction’s enzyme association can be factored into disjunctive normal form [22]. A rule in disjunctive normal form is a series of conjunctions (variables joined by *and*’s) connected by *or*’s into a disjunction. For enzyme associations, each conjunction describes a minimal set of enzymes or subunits that can independently catalyze a reaction. At least one conjunction of enzymes is necessary for a reaction to carry flux. In FALCON, complex enzyme associations are first factored into disjunctive normal form. The enzyme conjunctions are subsequently linked to reactions using a combination of the rules shown in Figure 2B and Figure 2D.

Reversible reactions must be split into two separate irreversible reactions – one carrying the forward flux and the other carrying the reverse flux (Figure 2E). Splitting reversible reactions ensures that enzyme activity is always consumed regardless of the flux direction. However, splitting reactions also creates the potential for a net consumption of enzyme activity with zero net flux through the associated reaction. If both the forward and reverse reactions carry the flux *v*, the net flux through the reaction will be *v − v* = 0 even though the reactions consume 2*v* of enzyme activity. To avoid these futile loops, we use a binary indicator variable to restrict flux through either the forward or reverse reactions. Using binary variable to restrict fluxes is a common approach to avoid infeasible loops in metabolic models [23]. The binary variables make FALCON models a non-convex MILP, as opposed to other convex LP formulations used to link enzymes and reactions [20, 21]. The added complexity of FALCON models is necessary to identify flux coupled reactions. If the binary variables are removed, any enzyme associated with a reversible reaction can decouple itself by forcing additional activity in the forward and reverse reactions. A complete description of the FALCON method is available in the Supplementary Materials.

**Table 1.** Model sizes for constraint-based reconstructions of *P. aeruginosa* [18], *S. mutans* [24], and *S. cerevisiae* before and after conversion to FALCON models.

| Model | Original Model |  |  | FALCON Model |  |
| --- | --- | --- | --- | --- | --- |
|  | Metabolites | Reactions | Enzymes | Constraints | Variables |
| *P. aeruginosa* | 1,300 | 1,551 | 1,148 | 6,696 | 6,946 |
| *S. mutans* | 511 | 715 | 400 | 1,907 | 2,105 |
| *S. cerevisiae* | 2,222 | 3,566 | 926 | 10,802 | 12,146 |

### Efficient flux coupling with solution caching

As shown in Figure 1, the coupling between enzymes is not the same as the coupling between reaction fluxes. The nonlinearities in the enzyme associations must be considered when searching for coupled enzyme sets. Since FALCON models include enzyme activities as variables in the metabolic model, we can in principle use Flux Coupling Finder (FCF) to identify coupled enzymes. Unfortunately, FALCON models are more difficult to solve due to their increased size and mixed-integer constraints. FCF requires solving 𝒪(*n*^2^) optimizations to identify coupled sets in a model with *n* variables [9]. Other methods exist to find coupled sets without solving optimizations [26, 9], but these methods require the model be convex (which mixed-integer models are not). Finding enzyme couplings with FALCON requires an efficient variant of FCF that supports non-convex mixed-integer models.

We developed an algorithm that uses caching to reduce the number of optimizations required by FCF. Our method, called cachedFCF, begins with the same strategy as FCF. We fix a single variable in the model (*x*_fixed_) to a nonzero value. We choose another variable *x*_*i*_ and perform two optimizations to find the maximum and minimum values for *x*_*i*_ while holding *x*_fixed_ constant. The variables *x*_*i*_ and *x*_fixed_ are fully coupled if and only if max(*x*_*i*_) = min(*x*_*i*_), i.e. if fixing *x*_fixed_ also fixes the value of *x*_*i*_. If max(*x*_*i*_) ≠ min(*x*_*i*_), we know that *x*_*i*_ and *x*_fixed_ cannot be fully coupled. FCF tests all pairs of variables for coupling. FCF uses shortcuts to avoid unnecessary optimizations. Fully coupling is symmetric, so testing *x*_*i*_ and *x*_*j*_ avoids the need to test for full coupling between *x*_*j*_ and *x*_*i*_. Similarly, any variable placed in a coupled set can be skipped when testing subsequent variables. Using these shortcuts, the original FCF study reduced the number of optimizations to far below the (*n*^2^ − *n*)/2 optimizations required for a brute-force approach on a model with *n* reactions [9].

By saving solutions for every optimization, we can further improve FCF’s efficiency. After testing if *x*_*i*_ is coupled to *x*_fixed_, FCF moves on to the next variable in the model, *x*_*i*+1_. Before testing if *x*_*i*+1_ is fixed, we check the solutions from maximizing and minimizing *x*_*i*_. If the value of *x*_*i*+1_ changed in either of the previous solutions, we know that *x*_*i*+1_ cannot be coupled to *x*_fixed_. If the value of *x*_*i*+1_ is unchanged we still need to maximize and minimize *x*_*i*+1_ before declaring that *x*_*i*+1_ and *x*_fixed_ are coupled. Moving on to variable *x*_*i*+2_, we check if *x*_*i*+2_ was not fixed in all the solutions found for *x*_*i*_ and *x*_*i*+1_. If so, we know *x*_*i*+2_ is not coupled to *x*_fixed_ without maximizing or minimizing *x*_*i*+2_.

In general, any variable that takes more than one value in any solution where *x*_fixed_ is constant cannot be coupled to *x*_fixed_. By saving the solutions to any optimization with constant *x*_fixed_, we can identify several variables that cannot be coupled to *x*_fixed_. Using the cached solutions avoids many costly optimization when searching for variables coupled to *x*_fixed_. Note that we cannot used cached solution to infer that two variables are coupled; a non-constant variable *x*_*i*_ cannot be coupled to a constant *x*_*fixed*_, but the converse is not true unless we have tried forcing *x*_*i*_ to move by maximizing and minimizing it. Most pairs of variables in a model are not coupled, so quickly identifying uncoupled variables can still yield substantial savings. Our cachedFCF algorithm requires up to three-fold fewer optimizations than standard FCF (Table 2). As expected, cachedFCF also reduced the runtime required to find all fully coupled reactions (Table 2).

We call saving all solutions for a single fixed variable (*x*_fixed_) “local caching". After searching for variables coupled to *x*_fixed_, we remove the constraint holding *x*_fixed_ constant and fix the next variable in the model. We must clear the local cache since our cachedFCF methodology only holds when all solutions contain a single fixed variable. Although we cannot directly test for coupling using solutions from a previous local cache, there is still valuable information in these solutions. All fully coupled variables in the model are perfectly correlated, i.e. the value of one variable is a scalar multiple of another. If we look across a large number of solutions, any two variables whose values are not perfectly correlated (or anticorrelated) cannot be coupled. Instead of discarding solutions from the local cache, we move them to a global cache. Before testing any pair of variables for coupling by optimization, we calculated the correlation coefficient between the variables using the samples in the global cache. The pair is not coupled and the optimizations are skipped unless the variables are significantly correlated.

Global caching and correlation testing drastically reduces the runtime of cachedFCF (Table 2). To limit the memory requirements for the global cache, we set an upper limit on the number of saved solutions (4,000 for our simulations). After the global cache is full, the cache is updated by randomly replacing existing solutions with new ones. A complete description of the cachedFCF algorithm can be found in the Supplementary Materials.

**Table 2.**
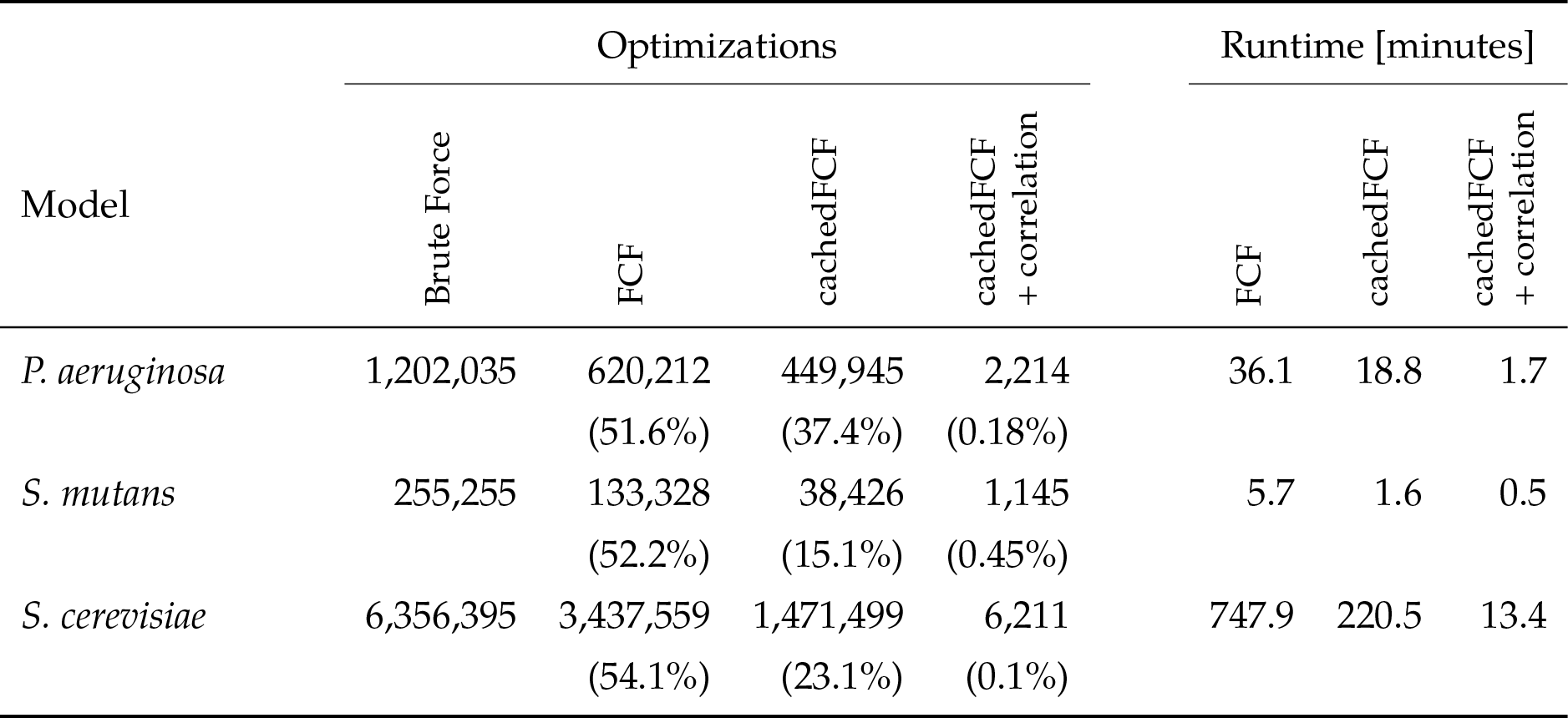
Caching reduces the number of optimizations and runtime when identifying coupled reaction sets. “Brute force” is the number of optimizations required to test all pairs without the shortcuts in FCF. “cachedFCF” uses only local caching. “cachedFCF + correlation” uses local and global caching. Analogous results for calculating coupled enzyme sets are presented in **Table S1**.

| Model | Optimizations |  |  |  | Runtime [minutes] |  |  |
| --- | --- | --- | --- | --- | --- | --- | --- |
|  | Brute Force | FCF | cachedFCF | cachedFCF + correlation | FCF | cachedFCF | cachedFCF + correlation |
| *P. aeruginosa* | 1,202,035 | 620,212<br>(51.6%) | 449,945<br>(37.4%) | 2,214<br>(0.18%) | 36.1 | 18.8 | 1.7 |
| *S. mutans* | 255,255 | 133,328<br>(52.2%) | 38,426<br>(15.1%) | 1,145<br>(0.45%) | 5.7 | 1.6 | 0.5 |
| *S. cerevisiae* | 6,356,395 | 3,437,559<br>(54.1%) | 1,471,499<br>(23.1%) | 6,211<br>(0.1%) | 747.9 | 220.5 | 13.4 |

### Enzymes in *ɛ*_FALCON_ sets are functionally coupled

We used cachedFCF to identify coupled enzyme sets in a genome-scale metabolic model of the human pathogen *Pseudomonas aeruginosa*. We calculated *E*_FCF_(*R*) sets by identifying fully coupled reaction sets and collecting enzymes associated with the reactions in each set. We also calculated *ɛ*_FALCON_ sets directly by converting the *P. aeruginosa* model to a FALCON model and applying cachedFCF. The *E*_FCF_(*R*) are larger (contain more enzymes) than the *ɛ*_FALCON_ sets calculated directly with the FALCON model (Figure 3a). By definition, enzymes belong to only one *ɛ*_FALCON_ set, whereas 38.9% (447/1149) of the enzymes in *P. aeruginosa* appear in more than one *E*_FCF_(*R*) set.

We tested if the enzymes in the smaller *ɛ*_FALCON_ sets show stronger functional connections. We previously used a collection of 214 gene expression datasets for *P. aeruginosa* to calculate the correlation coefficient for all pairs of metabolic genes [27]. Overall, pairs of genes assigned to the same *ɛ*_FALCON_ are more correlated than either pairs of genes selected from the entire metabolic network or the *E*(*R*) sets (Figure 3b). Each *E*_FCF_(*R*) set in *P. aeruginosa* contains one or more *ɛ*_FALCON_ set. We observed several cases where genes in the *ɛ*_FALCON_ sets are strongly correlated but genes in different *ɛ*_FALCON_ sets but the same *E*_FCF_(*R*) set are uncorrelated or anticorrelated (Figure 3c-d). Some, but not all, of the *ɛ*_FALCON_ sets are putative operons with adjacent genes on the genome. Genes in the larger *E*_FCF_(*R*) sets are often not near each other on the chromosome, which could contribute to the poor correlation between expression levels in some *E*_FCF_(*R*) sets.

Changes in gene expression or fitness assort non-randomly into enzyme sets. (We use the term “fitness change" to describe the change in a bacterium’s fitness when the gene’s function is interrupted.) If the *ɛ*_FALCON_ sets are fully coupled, we expect the genes in an *ɛ*_FALCON_ set to follow an “all-or-nothing" pattern regarding fitness or expression changes. If deleting one enzyme in a set creates a fitness defect, deleting any enzyme in the set should create a similar defect. We tested our “all-or-nothing" hypothesis using genome-wide expression (RNA-seq) and fitness (Tn-seq) profiles for *P. aeruginosa* during antibiotic stress [28] or a transition from *in vitro* minimal media to an *in vivo* mouse infection model [29]. We compared the fraction of *ɛ*_FALCON_ sets where all enzymes had significant expression changes or fitness changes. As a control, we randomly reassigned the fitness and expression changes to genes in *P. aeruginosa* (Figure 3c-d). Consistent with our hypothesis, the fraction of “all-or-nothing" sets was higher than expected; the fitness and expression changes cluster in the *ɛ*_FALCON_ sets.

**Figure 3.**
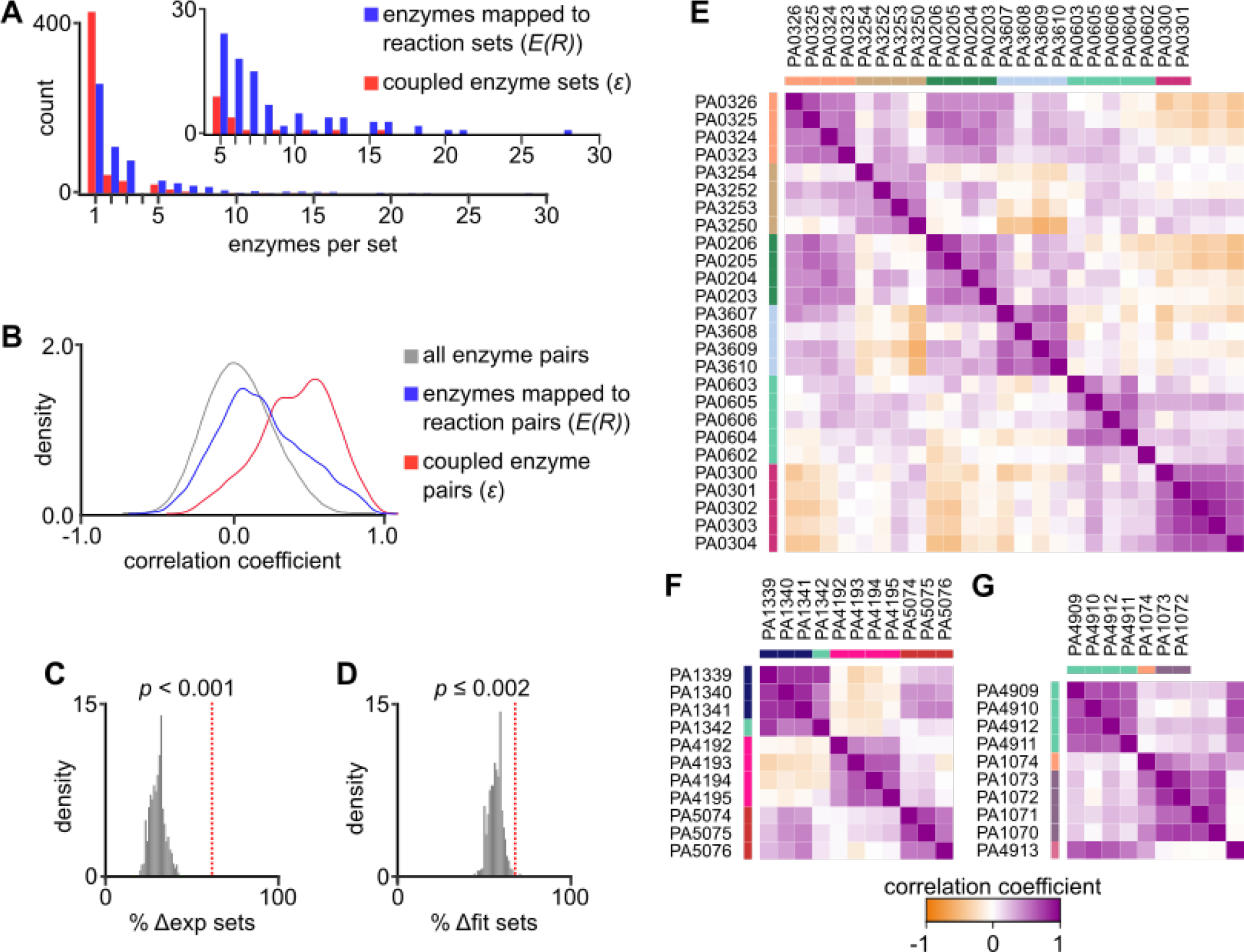
**A.** Enzyme sets computed directly from a FALCON model of *P. aeruginosa* are smaller than sets of enzymes associated with coupled reaction sets. The inset highlights sets with five or more enzymes per set. **B.** A meta-analysis of *P. aeruginosa* expression data [27] reveals strong pairwise correlations between enzymes in directly computed sets. The pairwise correlations are weaker among enzymes associated with the same coupled reaction set. **C.** Data from stress response studies in *P. aeruginosa* were used to test if gene expression or fitness changes clustered in enzyme sets. The observed number of gene sets with more than 50% of the enzyme differentially expressed (Δexp sets) is higher than a null distribution where the same number of expression changes were randomly assigned to genes. **D.** The number of sets with the majority of enzymes associated with a fitness defect (Δfit sets) is also higher than expected. In (**C**-**D**) the red dashed line marks the observed value in *P. aeruginosa*. The grey histograms summarize 1000 randomization experiments. **E-G.** Pairwise gene expression correlations in *E*_FCF_(*R*) sets. Each panel (**E**, **F**, or **G** shows genes in a single *E*_FCF_(*R*) set. The colored sidebars indicate the *ɛ*_FALCON_ sets inside the *E*(*R*) set. Genes in *ɛ*_FALCON_ sets are correlated, but genes in the same *E*_FCF_(*R*) set but different Efalcon sets are weakly or anti-correlated.

### Expanding *ɛ*_FALCON_ sets to branched and redundant pathways

Enzyme sets defined by flux coupling are used as surrogate pathways when analyzing high-throughput data. Most pathway databases or gene ontologies define pathways based on the metabolic network of a model organism, usually humans [12, 13]. Metabolic pathways vary widely across organisms, limiting the interpretability of pathways defined for other organisms. For example, the TCA cycle central is present in many organisms but absent in lactic acid bacteria. Some of the TCA enzymes are functional in lactic acid bacteria, but these enzymes are used as branchpoints into other pathways, not as a complete cycle [30]. Defining pathways *de novo* with enzyme sets avoids biasing our view of an organism’s metabolism with another organism’s pathways.

The enzyme sets identified by full flux coupling (and the weaker directional or partial coupling) are often incomplete when compared with standard pathways. Consider the example pathway in Figure 4a. In *P. aeruginosa*, the transformation of 5-dehydroshikimate to 3-dehydroquinate can be independently catalyzed by either enzyme PA4846 or enzyme PA0245. Most biologists and pathway databases consider this set of four reactions to be one pathway. However, only two of the reactions (the reactions not catalyzed by the isozymes) are fully coupled by flux coupling analysis. Depending on the reversibility of the reactions, all four reactions are not always directionally or partially coupled either.

The redundancy in a four reaction, branched pathway (Figure 4b) prevents coupling of the entire pathway. Since flux can can travel through either branch, the branched reactions will never be coupled to the unbranched reactions. However, if one of the branched reactions is removed, the remaining three reactions are fully coupled since flux must travel through the remaining branch. We observed that any singly branched pathway can be forced into a subset of fully coupled reactions by removing one of the branches. (For pathways with three or more branches, all but one of the branches need to be removed to force coupling.) Based on this observation, we developed a computational method called “delete-and-couple” to identify coupled sets from redundant or branched pathways. Delete-and-couple removes one reaction from the model and identifies fully coupled sets using cachedFCF. The process is repeated by removing every reaction individually. The coupled sets from all the deletions are pooled and merged based on shared reactions. For the simple branched pathway in Figure 4d, single reaction deletions create two coupled sets containing either the left or right branch. Because both sets share the unbranched reactions, they are merged into a single, branched pathway.

We ran our delete-and-couple algorithm on the *P. aeruginosa* genome-scale model. For efficiency, we deleted only one reaction in each coupled reaction set since deleting any reaction in a coupled set has the same effect on the model. Fortunately, the efficiency of the cachedFCF makes delete-and-couple tractable, requiring 1.7 minutes to complete on a desktop computer. We found “expanded" enzyme sets for *P. aeruginosa* in two ways. First, we applied delete-and-couple to the *P. aeruginosa* metabolic model to identify expanded reaction sets. We collected enzymes associated with the reactions in each set, producing *E*_FCF_(*R*\*) sets. (The *R*\* notation indicates that the reaction sets were expanded with delete-and-couple and subsequently mapped to enzymes.) We also calculated expanded enzyme sets directly (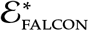 sets) by applying delete-and-couple to enzymes in a FALCON model. The 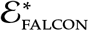 sets directly calculated with the FALCON model are smaller than the *E*_FCF_(*R*\*) sets, just as the *ɛ*_FALCON_ sets are smaller than the *E*_FCF_(*R*) sets (Figure 4e). On average, both the *E*_FCF_(*R*\*) and 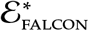 sets are larger than the *E*_FCF_(*R*) and *ɛ*_FALCON_ sets, respectively. The delete-and-couple 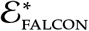 sets each contain one or more *ɛ*_FALCON_ sets. Visually, the expanded 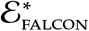 sets align with traditional views of metabolic pathways, while the *ɛ*_FALCON_ sets alone are often small linear branches of more complex pathways (Figure 5).

**Figure 4.**
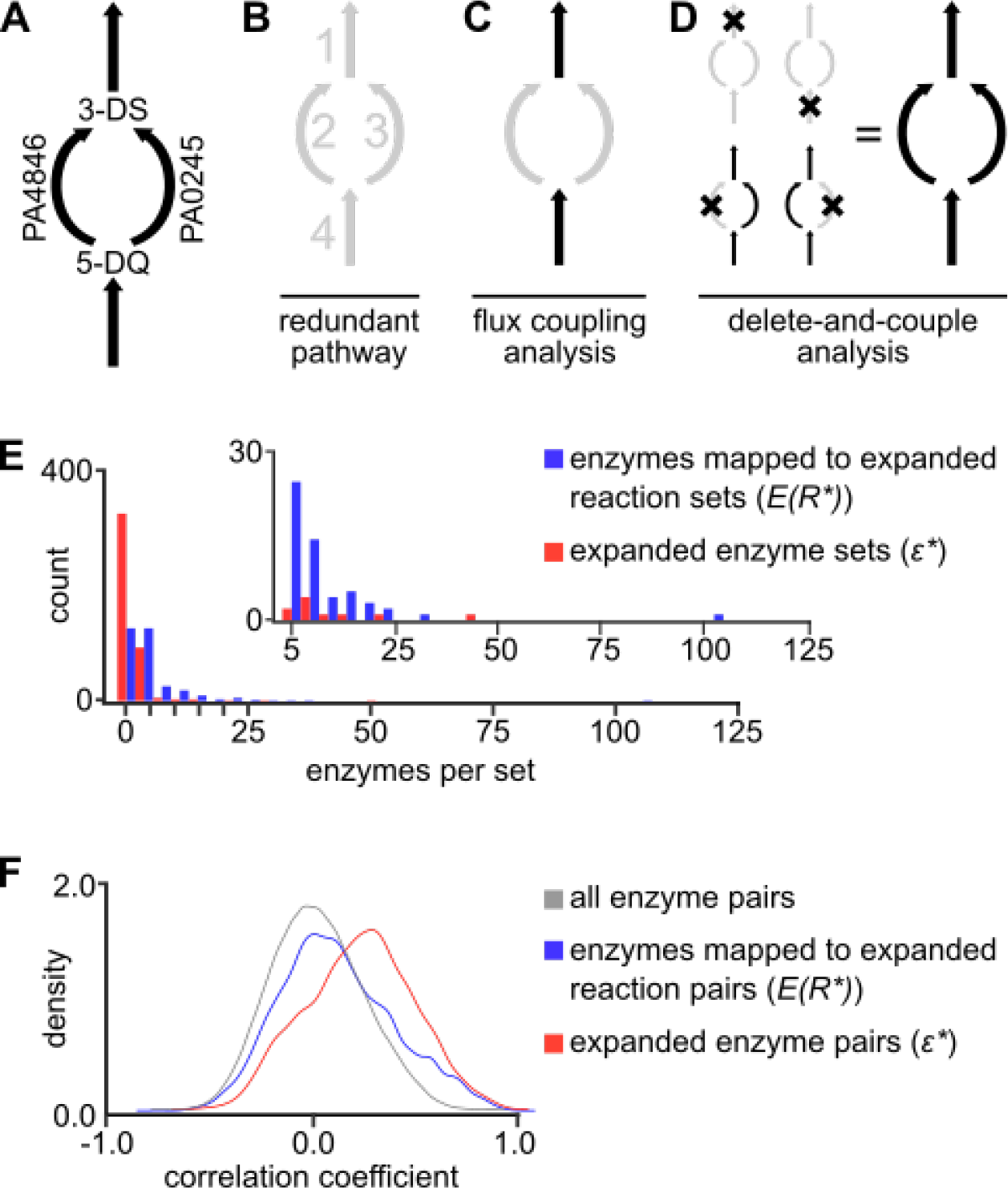
Branched pathways are not captured by flux coupling analysis. **A.** As an example branched pathway, either of two isozymes (PA4846 or PA0245) can independently catalyze the formation of 3-dehydroquinate (3-DS) from 5-dehydroshikimate (5-DQ). The redundancy in reactions 2 and 3 (**B**) are not coupled to either reaction 1 or reaction 4 by flux coupling analysis (**C**). **D**. If either of the branched reactions are removed, the remaining three reactions become fully coupled. The coupled sets from each perturbation can be merged into a complete pathway by delete-and-couple analysis. **E.** Expanded enzyme sets computed by delete-and-couple with a FALCON model are smaller than sets of enzymes mapped to expanded reaction sets. The inset highlights sets with five or more enzymes per set. **F.** Pairs of enzymes in the expanded enzyme sets retain pairwise correlation in *P. aeruginosa*, while enzymes mapped to expanded reaction sets do not.

**Figure 5.**
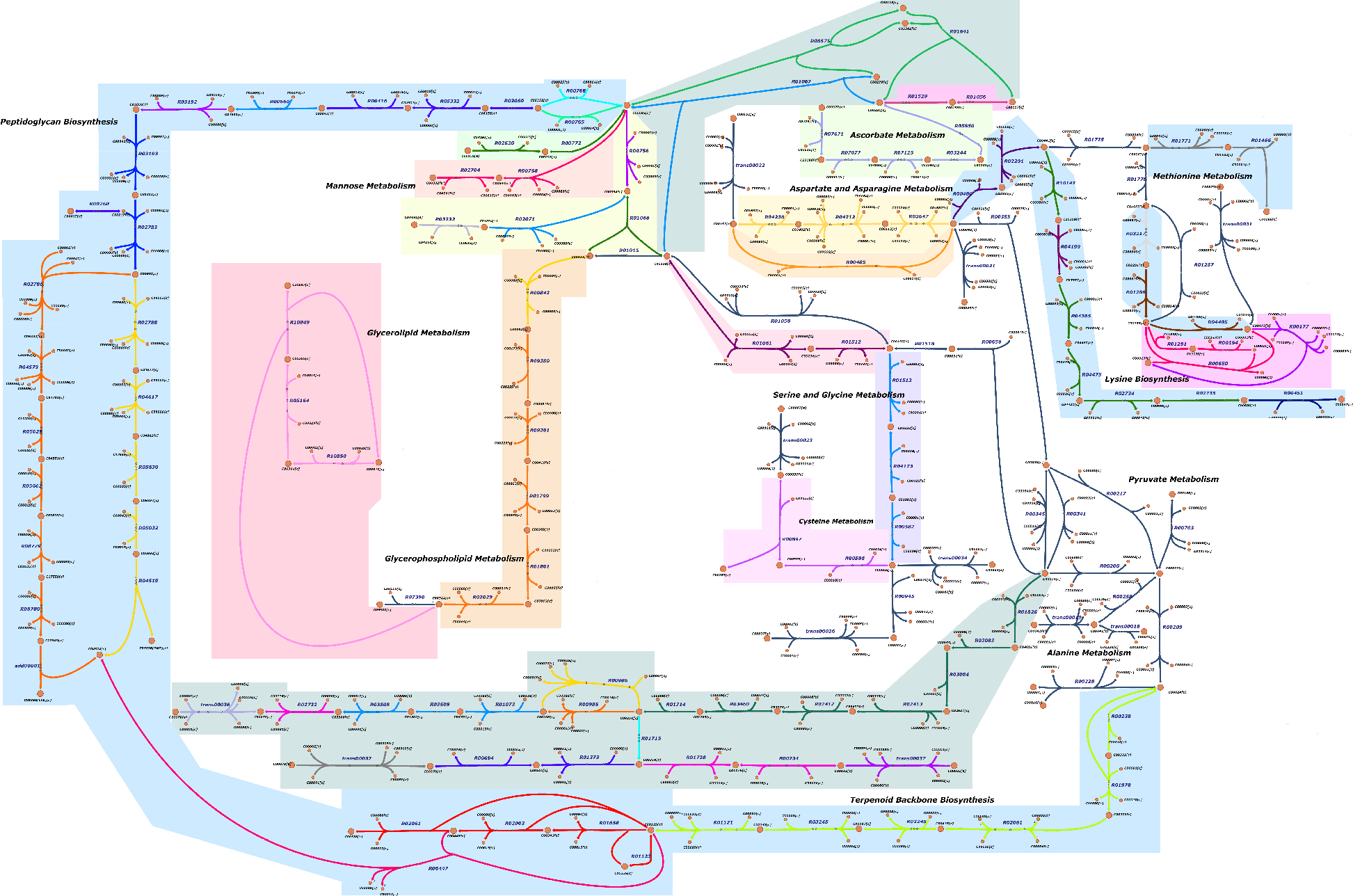
Expanded enzyme sets combine multiple enzyme sets into larger pathways. Individual enzyme sets (*ɛ*_FALCON_) for *S. mutans* (model iSMUv1 [24]) are shown by coloring the associated reactions. The expanded enzyme sets 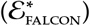 are represented by colored shading around the reactions.

The expanded 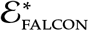 sets contain more enzymes per set, but the enzymes in each set remain functionally linked. Looking again at pairwise correlations in *P. aeruginosa* gene expression, we see that pairs of enzymes in the same 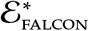 have higher gene expression correlation than randomly selected pairs of enzymes or enzymes in *E*_FCF_(*R*\*) sets (Figure 4f). The enzymes added when *ɛ*_FALCON_ sets are expanded with delete-and-couple are often co-expressed with other enzymes in the set.

Our analysis also showed that fitness and expression changes cluster within *ɛ*_FALCON_ sets. Multiple *ɛ*_FALCON_ sets comprise each 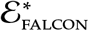 sets, but are the *ɛ*_FALCON_ sets in each 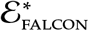 alike? Do the *ɛ*_FALCON_ sets with all expression changes join together, or are fitness and expression changes consolidated into 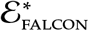 sets? We can view each 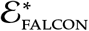 set as a network of connected *ɛ*_FALCON_ sets (Figure 6a). The *ɛ*_FALCON_ sets are connected based on shared enzymes identified during the delete-and-couple procedure. Two *ɛ*_FALCON_ sets are connected inside a 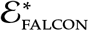 set if the *ɛ*_FALCON_ sets were fully coupled during at least one perturbation. We tested how often delete-and-couple connected pairs of *ɛ*_FALCON_ sets with fitness or expression changes. Sets with expression changes (Δexp sets) connected with other Δexp sets more often than expected by chance (Figure 6b). Sets with fitness changes (Δfit sets) connected with other Δfit sets more often than expected. We also observed frequent connections involving both a sets with fitness and expression changes (Figure 6c). Of all the ways to connect sets with both fitness and expression changes, only the case where a set with both expression and fitness changes connects to another set with both changes is overrepresented (Figure 6e-g). Other cases, such as connections between a Δfit set and a Δexp set (Figure 6e), occur randomly.

### Implementation

The cachedFCF, FALCON, and delete-and-couple algorithms were written in R using the sybil package [31]. All source code (including a Python implementation of cachedFCF) is freely available at the authors’ website: http://jensenlab.net/tools. Simulations were performed on a quad-core 3.2 GHz Intel i7 workstation with 48 GB of RAM using Gurobi version 8.0.

## Discussion

Flux coupling is a powerful tool for analyzing genome-scale, constraint-based models. Unfortunately, users of current flux coupling algorithms face three challenges: 1.) identifying coupled reactions is computationally expensive, especially for non-convex models; 2.) complex associations between enzymes and reactions make it difficult to translate coupled reaction sets into coupled enzyme sets; and 3.) fully coupled sets are often too small, while partial or directionally coupled sets can be large and diffuse [9]. Our cachedFCF, FALCON, and delete-and-couple tools overcome these obstacles.

Like the original FCF algorithm, cachedFCF has a worst-case runtime that scales quadratically with the number of model variables. In practice, we see a 100-1000 fold reduction in the number of optimizations and a 10-100 fold decrease in runtime using cachedFCF. The efficiency of cachedFCF enables new applications of flux coupling, such as our delete-and-couple method. Delete-and-couple has cubic worst-case scaling but runs in hours for the *P. aeruginosa* genome-scale model.

**Figure 6.**
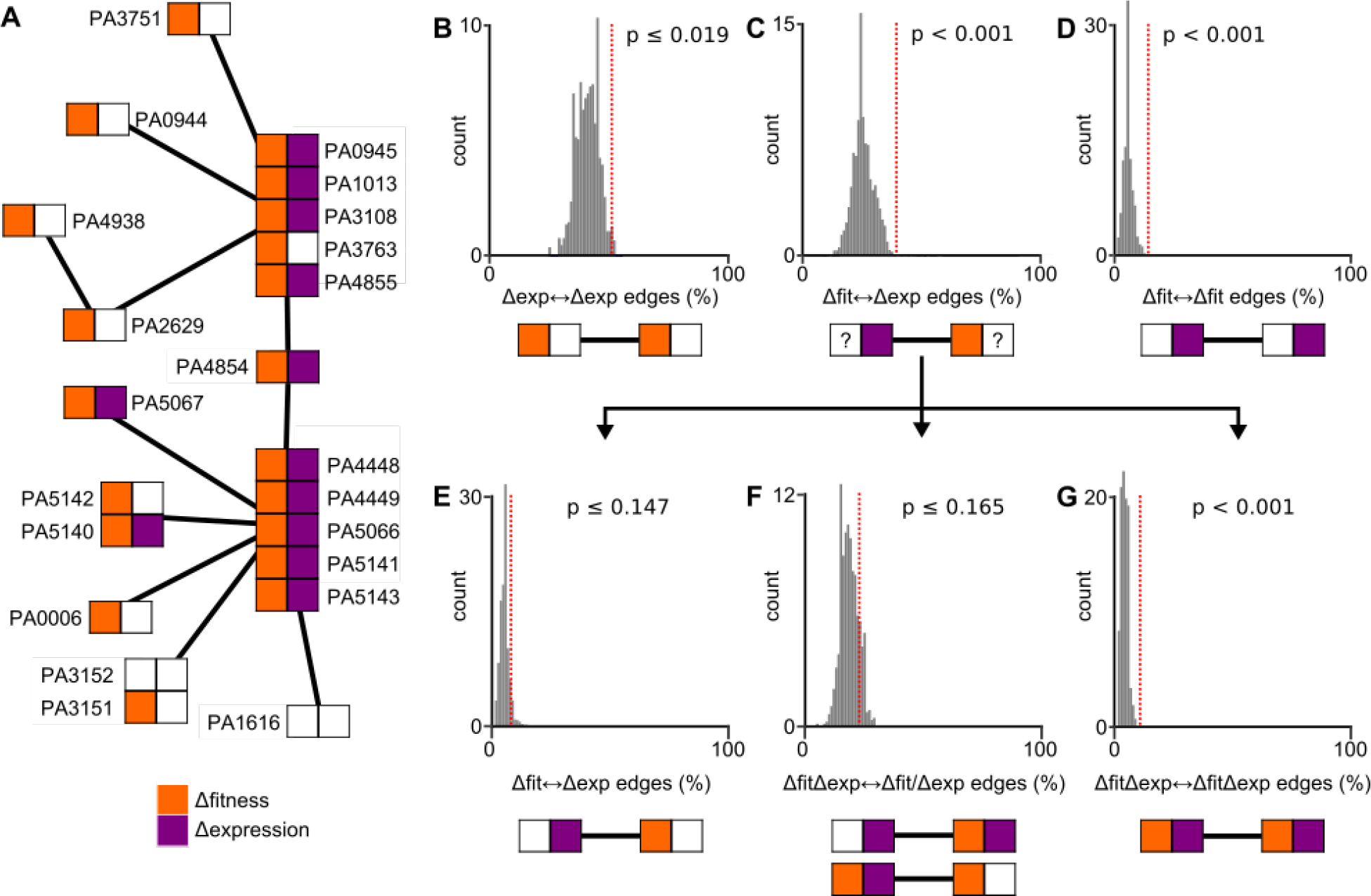
Expanded enzyme sets 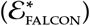 link enzyme sets with similar functional profiles. **A.** An expanded enzyme set viewed as a network of individual enzymes sets (*ɛ*_FALCON_). Two *ɛ*_FALCON_ sets are connected if they are joined by shared enzymes during a perturbation in the delete-and-couple analysis. To test if *ɛ*_FALCON_ sets join with similar *ɛ*_FALCON_ sets, we compared the frequency of connections in *P. aeruginosa* (red dashed line) with a null distribution of 1000 simulations where Δexp and Δfit changes are randomly assigned to genes (grey histograms). Connections between two Δexp sets or two Δfit sets occur more frequently than expected (**B**, **D**). Connections between two sets that both have Δexp and Δfit genes are also overrepresented (**G**). Other connections (**E**, **F**) occur randomly.

The COBRA Toolbox [14] uses random sampling to identify flux couplings by calculating correlation coefficients between all pairs of fluxes. Our global cache screening takes a similar approach, although we use the lack of correlation to identify non-coupled reactions. By only skipping pairs of reactions with clear evidence of no coupling, cachedFCF avoids missing couplings due to incomplete or non-uniform random sampling. We believe cachedFCF balances the efficiency of random sampling with the completeness of the original FCF algorithm. Most importantly, cachedFCF makes no assumptions about the structure of the metabolic model. The algorithm is compatible with any mathematical program (LPs, MILPs, MIQCPs, etc.), allowing researchers to add regulatory constraints or true enzyme associations with FALCON.

FALCON is not the first framework to incorporate enzyme activities as continuous variables in constraint-based models [20, 21]. It is the first approach that fully couples activity to the corresponding reaction fluxes. FALCON models require binary variables to prevent cycles of activity in reversible reactions. FALCON models are non-convex because of the binary variables. Activity cycles are not a problem for many FBA-type analyses where surplus enzyme activity would be suboptimal. For algorithms like FCF, activity cycles can break the correlation between coupled reactions by allowing physiologically meaningless activity distributions. While FALCON models are more computationally demanding than models using other frameworks, they allow analyses like flux coupling to be applied directly to enzymes.

Using FALCON models, many constraint-based modeling algorithms that operate on reactions can be applied to enzymes. Combining FALCON with common methods like flux variability analysis [32], MOMA [33], and random sampling [34] could offer new insights on how the enzymatic network coordinates the metabolic network. Gene or protein expression profiles are far cheaper and more comprehensive than flux profiles, so moving to an enzyme-centric view of metabolism would accelerate data-driven approaches to constraint-based modeling.

## Acknowledgements

We thank Kevin D’Auria and Matt Biggs for feedback on the FALCON framework. This work was supported by the National Institutes of Health (grant EB027396 to PJ and GM088244 to JP) and a National Science Foundation graduate fellowship to PJ. The authors declare no financial or commercial conflict of interest.

