## Supplementary Material for "Efficient enzyme coupling algorithms identify functional pathways in genome-scale metabolic models"

**Table S1.** Caching reduces the number of optimizations and runtime when identifying coupled enzyme sets. “Brute force" is the number of optimizations required to test all pairs without the shortcuts in FCF. “cachedFCF" uses only local caching. “cachedFCF + correlation" uses local and global caching.

| | Optimizations | | | | Runtime [minutes] | | |
|---|---|---|---|---|---|---|---|
| Model | Brute Force | FCF | cachedFCF | cachedFCF + correlation | FCF | cachedFCF | cachedFCF + correlation |
| *P. aeruginosa* | 1,010,331 | 630,213 | 385,690 | 2,285 | 304.1 | 176.1 | 4.3 |
| | | (62.4%) | (38.1%) | (0.2%) | | | |
| *S. mutans* | 193,131 | 115,283 | 44,599 | 1,096 | 10.8 | 0.8 | 0.7 |
| | | (59.7%) | (23.1%) | (0.6%) | | | |
| *S. cerevisiae* | 11,589,705 | 1,558,818 | 868,876 | 4,594 | 1,789.6 | 927.8 | 15.5 |
| | | (13.4%) | (7.5%) | (0.04%) | | | |

**Data:** MILP COBRA model
**Result:** Boolean matrix indicating coupling between variables
$N_{\text{vars}}$= number of variables in model;
$c_{\text{thresh}}$ = threshold for correlation between variables;
$x^{\text{global}}_{\text{max}}[N_{\text{vars}}]$ = maximum observed values for each variable. Initialized to 0;
$x^{\text{global}}_{\text{max}}[N_{\text{vars}}]$ = minimum observed values for each variable. Initialized to 0;
$x^{\text{local}}_{\text{max}}[N_{\text{vars}}]$ = maximum observed values when a single variable is fixed. Initialized to Infinity;
$x^{\text{local}}_{\text{min}}[N_{\text{vars}}]$ = minimum observed values when a single variable is fixed. Initialized to -Infinity;
Cache[$N_{\text{vars}}$,$N_{\text{cache}}$] = matrix of cached solutions. Initialized to 0;
**Routine** update(solution[$N_{\text{vars}}$]) **:**
  **for** $i \in 1 : N_{\text{vars}}$ **do**
    $x^{\text{global}}_{\text{min}}[i] = \min\{x^{\text{global}}_{\text{min}}[i], \text{solution}[i]\}$;
    $x^{\text{global}}_{\text{max}}[i] = \max\{x^{\text{global}}_{\text{max}}[i], \text{solution}[i]\}$;
    $x^{\text{local}}_{\text{min}}[i] = \min\{x^{\text{local}}_{\text{min}}[i], \text{solution}[i]\}$;
    $x^{\text{local}}_{\text{max}}[i] = \max\{x^{\text{local}}_{\text{max}}[i], \text{solution}[i]\}$;
    **if** Cache is full **then**
      add solution to Cache;
    **else**
      replace oldest Cache entry with new solution;

Coupled[$N_{\text{vars}}$,$N_{\text{vars}}$] = coupling matrix. If Coupled[$i$,$j$] is true, variables $i$ and $j$ are coupled;
isBlocked[$N_{\text{vars}}$] = vector identifying blocked variables. Initialized to false;
isCoupled[$N_{\text{vars}}$] = vector identifying variables that are not yet coupled. Initialized to false;
**for** $i$ in model.variables **do**
  reinitialize $x^{\text{local}}_{\text{min}}$ and $x^{\text{local}}_{\text{max}}$;
  **if** model.ub[$i$] $\neq 0$ **then**
    solution ← maximize variable $i$;
    **update**(solution);
  **if** model.lb[$i$] $\neq 0$ **then**
    solution ← minimize variable $i$;
    **update**(solution);
  **if** $x^{\text{global}}_{\text{max}}[i] = x^{\text{global}}_{\text{min}}[i] = 0$ **then**
    isBlocked[$i$] = true;
  **else**
    Coupled[$i$,$i$] = true;
  isCoupled[$i$] = false;
  choose $\epsilon \neq 0 \in (x^{\text{global}}_{\text{min}}[i], x^{\text{global}}_{\text{max}}[i])$;
  fix variable $i$ at $\epsilon$ (model.lb[$i$] = $\epsilon$ and model.ub[$i$] = $\epsilon$);
  **for** $j$ in model.variables[$i+1$:end] **do**
    **if** isBlocked[$j$] or isCoupled[$j$] **then** next $j$;
    **if** $x^{\text{local}}_{\text{max}} \neq x^{\text{local}}_{\text{min}}$ **then** next $j$;
    **if** correlation(Cache[:,$i$], Cache[:,$j$]) $\leq c_{\text{thresh}}$ **then** next $j$;
    solution ← maximize variable $j$;
    **update**(solution);
    **if** $x^{\text{local}}_{\text{max}} = x^{\text{local}}_{\text{min}}$ **then**
      solution ← minimize variable $j$;
      **update**(solution);
    **if** $x^{\text{local}}_{\text{max}} = x^{\text{local}}_{\text{min}}$ **then**
      Coupled[$i$,$j$] = true;
      isCoupled[$j$] = true;
  reset model.lb[$i$] and model.ub[$i$];
return Coupled;

**Algorithm 1:** Cached Flux Coupling Finder (cachedFCF).

### Converting COBRA models into FALCON models.

We define three classes of reactions based on the degeneracy of their enzyme associations.

- **Class Ia** reactions are associated with enzymes that are exclusive to the reaction – the enzymes are not associated with any other reaction. Additionally, the DNF representation of the GPR rule has only a single clause (see below). This is equivalent to having no *or*'s in the GPR rule.
- **Class Ib** reactions also associated with enzymes that are exclusive to the reaction. Unlike Class Ia reactions, the DNF representation of the GPR rules for Class Ib reactions can contain multiple clauses.
- **Class II** are associated with enzymes that are also associated with another (Class II) reaction.

Adding enzyme associations requires the boolean GPR rules be factored into the disjunctive normal form (DNF). A DNF rule is a series of one or more *clauses* joined by *or*'s. Each clause contains one or more enzyme joined by *and*'s. For example, the GPR rule

$$e_1 \text{ and } (e_2 \text{ or } e_3)$$

is not a DNF rule. It can be factored into the equivalent DNF rule

$$(e_1 \text{ and } e_2) \text{ or } (e_1 \text{ and } e_3)$$

For GPR rules, each clause in the DNF represents a set of enzymes that are sufficient to catalyze the enzyme. FALCON creates a separate reaction for each clause in the DNF. These reactions are linked to enforce mass balance constraints and prevent futile cycles. Algorithm 2 presents the complete FALCON method for linking enzyme activities to reactions in COBRA models.

```
Data: LP COBRA model
Result: MILP FALCON model
Routine coupleEnzymesToReaction(reaction, enzymes, reversible) :
    if reversible then
        add enzymes as pseudo-reactants to reaction;
    else
        split reaction into forward and reverse reactions;
        add enzymes as pseudo-reactants to both reactions;
Routine addEnzymeClausesToReaction(reaction, clauses, reversible, split) :
    add new metabolite “rxnActivity" to model;
    for clause in clauses do
        if split then
            add two reactions (forward and reverse) that consume enzymes in clause and produce rxnActicity;
        else
            add one reversible reaction that consumes enzymes in clause and produces rxnActivity;
        addEnzymesToReaction(reaction, enzymes=rxnActivity, reversible=reversible);
    add indicator variable so only forward or reverse reactions can carry flux not both;
for reaction in Class Ia do
    addEnzymesToReaction(reaction, enzymes(reaction), split=false);
for reaction in Class Ib do
    addEnzymesClausesToReaction(reaction, DNF(enzymes(reaction)), reversible=true, split=true);
for reaction in Class II do
    addEnzymesClausesToReaction(reaction, DNF(enzymes(reaction)), reversible=false, split=true);
```

**Algorithm 2:** Creating a FALCON model.
